# Visual modelling supports the potential for prey detection by means of diurnal active photolocation in a small cryptobenthic fish

**DOI:** 10.1101/338640

**Authors:** Pierre-Paul Bitton, Sebastian Alejandro Yun Christmann, Matteo Santon, Ulrike K. Harant, Nico K. Michiels

## Abstract

Active sensing has been well documented in animals that use echolocation and electrolocation. Active photolocation, or active sensing using light, has received much less attention, and only in bioluminescent nocturnal species. However, evidence has suggested the diurnal triplefin *Tripterygion delaisi* uses controlled iris radiance, termed ocular sparks, for prey detection. While this form of diurnal active photolocation was behaviourally described, a study exploring the physical process would provide compelling support for this mechanism. In this paper, we investigate the conditions under which diurnal active photolocation could assist *T. delaisi* in detecting potential prey. In the field, we sampled gammarids (genus *Cheirocratus*) and characterized the spectral properties of their eyes, which possess strong directional reflectors. In the laboratory, we quantified ocular sparks size and their angle-dependent radiance. Combined with environmental light measurements and known properties of the visual system of *T. delaisi*, we modeled diurnal active photolocation under various scenarios. Our results corroborate that diurnal active photolocation should help *T. deia¡s¡* detect gammarids at distances relevant to foraging, 4.5 cm under favourable conditions and up to 2.5 cm under average conditions. Because ocular sparks are widespread across fish species, diurnal active photolocation for micro-prey may be a common predation strategy.

## Introduction

Active sensory systems have been well studied in several animals. For example, the echolocating behavior of bats, by which the reflection of emitted sound waves contributes to navigation in the dark, has been detailed since 1938^1,2^, and active electrolocation, by which the disruptions of weak electrical fields are used to detect potential prey and predators, is well known from model organisms such as the electric fish *Apteronotus leptorhynchus*^3,4^. In contrast, active photolocation, the process by which organisms actively use light to survey their environment, seems limited to bioluminescent organisms; only deep-sea dragonfish (family Stomiidae), lanternfish (family Myctidae), and nocturnal flashlight fish (family Anomalopidae) are assumed to use active photolocation^5–7^. However, recent evidence suggests that active photolocation, by controlled light redirection, could also be used in diurnal fish to assist in prey detection, and may be generally common across fish species^8^.

Michiels et al.^8^ described a mechanism that allows the triplefin *Tripterygion delaisi* to redirect ambient light by taking advantage of its laterally protruding lenses and reflective irides, and discussed how this may assist in the detection of camouflaged micro-prey. The central basis of the mechanism is that downwelling light strikes the dorsal part of the eye, is focused by the protruding lens onto the iris below the pupil, and is reflected in the horizontal plane of vision. The focussed light can be radiated by the red fluorescent section of the iris producing a ‘red ocular spark’, reflected by a blue-white area below the pupil generating a ‘blue ocular spark’ (**Fig. 1**), or turned on and off by rotating and tilting the iris (see **Fig. 2** in ^8^). Their experiments demonstrated that the absence/presence and type of sparks (red and blue) depends on the environmental context, and is under voluntary control^8^. Because downwelling light in the aquatic environment is many times more intense than sidewelling light^9,10^, blue ocular sparks are brighter than the background. Michiels et al.^8^ emphasized that ocular sparks are too weak to illuminate an entire scene, but suggested they may be sufficiently radiant to reveal strong and/or directional reflectors in nearby target organisms.

**Figure 1.**
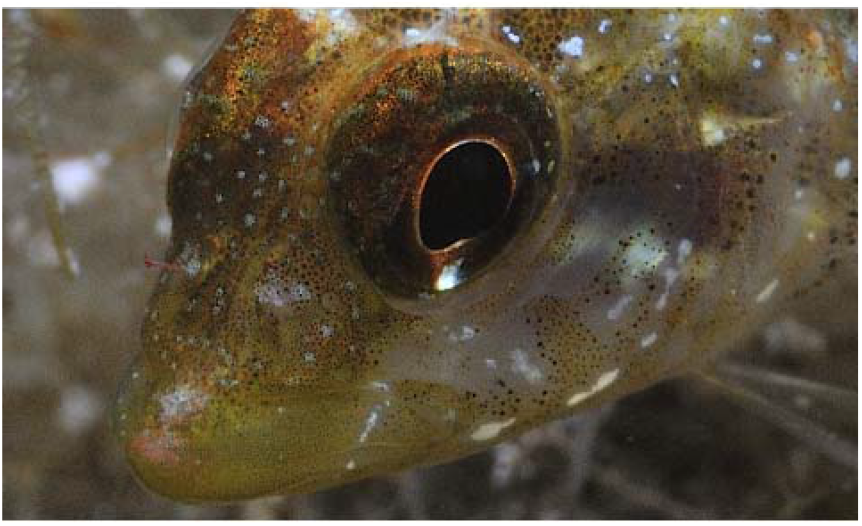
The triplefin *Tripterygion delaisi* produces blue ocular sparks (bright area ventral to the pupil) by focussing downwelling light onto blue chromatophores. This downwelling light is reflected equally in all directions (See **Fig. 2**). Photo credit: Nico K. Michiels

Indeed, strongly reflecting structures are abundant in aquatic ecosystems, specifically in the eyes of both vertebrates and invertebrates^11,14^. For example, camera eyes that possess either a *tapetum lucidum* or *stratum argenteum* are retroreflective, and produce the eyeshine observed when illuminating nocturnal animals. This type of reflected eyeshine is only perceived if the illuminating source is coaxial to the receiver’s eye because most of the light is returned to the source in a narrow angle. Furthermore, invertebrates such as stomatopod larvae also possess strong reflectors considered to camouflage their opaque retinas^12^. Though not true retroreflectors, the reflectance of marine invertebrate compound eyes is often stronger with a coincident normal viewing geometry i.e., coaxial alignment^8,12,15^. Strong directional reflectors and coaxially generated illumination are key components of the mechanism proposed by Michiels et al.^8^ because the ocular sparks are produced on the irides, immediately adjacent to the pupil. Thus, ocular sparks could make use of the reflectance of prey eyes to increase the probability of detection, as has been suggested for nocturnal, bioluminescent species^15–17^.

The experiment reported in Michiels et al.^8^ was conducted in the laboratory and focused on ocular spark modulation in response to prey presence and background hue. No studies have yet explored the physical and theoretical basis of the complete process to describe the conditions under which ocular sparks could assist triplefins in detecting prey in natural contexts. In this study, we use simple mathematical expressions and visual modelling to determine the conditions that would enable triplefins to benefit from blue ocular sparks for prey detection. In the field, we collected measurements of ambient light and characterised the reflective properties of a background in which gammarids (Crustacea: Amphipoda), important triplefin prey items^18,19^, are found. In the laboratory, we measured the ocular spark properties of *T. delaisi* and the optical properties of the eyes and bodies of gammarids. Finally, we combined these data with *T. delaisi’s* species-specific visual system characteristics^20^ to inform models of visual interactions between *T. delaisi* and gammarids while varying influential parameters.

## Materials and Methods

The yellow black-faced triplefin *Tripterygion delaisi* is a small benthic fish found along Mediterranean and eastern Atlantic coasts^21^. It lives in rocky habitats between 5 and 50 m depth where it feeds mainly on small benthic invertebrates^18,22^. We study them at the Station de Recherches Sous-marines et Océanographiques (STARESO) in Calvi (Corsica, France), where our preliminary investigations suggest that their preferred food items include gammarids, caprellids, copepods, and decapods (Fritsch and Michiels, unpublished data). Microscopy investigations have shown that almost all of these possess strong reflectors in their eyes (Bitton, Fritsch and Michiels, unpublished data), which concurs with the existing literature on aquatic invertebrates^11,12,23-27^. Triplefins are highly cryptic against their natural background with no obvious sexual dimorphism, except during the breeding season when males acquire dark heads and yellow bodies.

### Field light environments and background reflectance

We measured the reflective properties of *Halopteris filicina*, a common foraging substrate for *T. delaisi*, and the downwelling light, unshaded sidewelling light, and shaded sidewelling light of triplefin habitat at STARESO in June-July 2014 and 2017. Details of *Halopteris filicina* data collection protocol can be found in Harant et al.^19^ In short, substrate data were collected while scuba diving at a shallow site (5 m) characterized by rocky slopes, steep walls and granite boulders. Measurements were obtained at various locations in conjunction with a polytetrafluorethylen (PTFE) diffuse white standard (DWS; Berghof Fluoroplastic Technology GmbH, Eningen unter Achalm, Germany) tilted at 45° to the surface as a combined measure of downwelling and sidewelling light. The relative radiance between the substrate measurements and light field is considered as the reflective property of *Halopteris filicina*. Light field measurements were obtained between 2 and 30 m depth on substrates facing south. At each depth (2, 4, 6, 8, 10, 14, 18, 24, and 30 m) we measured from a 45° angle the radiance of an exposed PTFE standard set at normal incidence to the water surface (= angle of incidence 0°) for an approximation of downwelling light, from a PTFE standard set at 90° to normal for measuring sidewelling light, and a PTFE standard set at 90° to normal and shaded by a 4 cm opaque black cover as a measure of shaded sidewelling light environment. Three measurements were obtained for every standard at every depth and averages used in analyses. All measurements were obtained using a SpectraScan^®^ PR-740 (PhotoResearch Inc., Syracuse, USA) fixed at a focal distance of 50 cm in a custom-built underwater housing (BS Kinetics, Achern, Germany). The SpectraScan spectroradiometers use Pritchard optics to collect measurements of absolute radiance from a specific solid angle, which is visualized as a small black circular area in the viewfinder. The PR-740 was equipped with a colour correction filter (#287 double CT orange, LEE Filters, Andover, England) which suppresses but does not block the dominant blue-green spectral range. This increases exposure time, allowing the instrument to obtain better readings in the weak, long-wavelength part of the spectrum at depth. Radiance measurements were corrected for the transmission profile of the filter and port of the housing before being used in the calculations.

### Properties of gammarids

We isolated gammarids from *Halopteris filicina* algae collected between 5 and 10 m depth at STARESO, and kept them immobilized but alive using a 0.6 M MgCl_2_ solution. Spectral measurements of their body and compound eye were obtained with a PR-740 spectroradiometer mounted onto a Leica DM5000 B compound microscope (Leica Microsystems, Wetzlar, Germany) under 10 × 10 magnification. For reflectance measurements, we used an external halogen light source (KL2500 LCD, Schott AG, Mainz, Germany), either coincident through the microscope’s housing (epi-illumination) or at 45° to the sample using an external LLG 380 liquid light guide (Lumatec GMBH, Germany). For each gammarid we collected five body and eye reflectance measurements coaxially illuminated, and five measurements of eyes illuminated at 45° relative to the optical axis. We did not collect body reflectance at 45° because there was no evidence of specularity or iridescence. The measurement area covered almost the entire gammarid eye under 10 x 10 magnification. A submerged PTFE standard was also measured under comparable geometries five times both with coaxial epi-illumination and with the light source at 45°. In all cases, the sample was repositioned and refocused before each measurement. Averages of 5 measurements of the body and eyes were expressed in relation to their relative standard. For transmission measurements we used the 12 V 100 W halogen lamp provided with the microscope in the transmitted light axis. For each gammarid, we took five radiance measurements of the transmitted light as seen through haphazardly selected locations on the body (plus Petri dish and MgCl_2_ solution) and five reference measurements of the transmitted light without the gammarid (but including the Petri dish and MgCl_2_ solution). Transmittance was then determined as the mean of the five measurements of the body divided by the reference. Scaled images of the gammarids (full length 3 – 4 mm) were also obtained at this time and the size of the eyes subsequently measured using ImageJ^28^.

### Properties of ocular sparks

Radiance of the blue ocular spark was measured in live fish and the results are reported in Michiels et al.^8^. In short, individual fish (*n* = 5) were held in a small tank (L⍰×⍰W⍰×⍰H⍰=10⍰×⍰6⍰×⍰4 cm^3^) in a dark room, illuminated from above through means of a liquid light guide attached to a broad-spectrum EL 6000 source (Leica Microsystems, IL USA). The radiances of sparks and a Spectralon diffuse white reflectance standard (SRS-99-010; Labsphere Inc, NH USA) were measured with a calibrated SpectraScan^®^ PR-670 spectroradiometer (PhotoResearch Inc, NY USA). The reflectance of the spark was calculated as the quotient of the spark radiance divided by the radiance of the white standard for every fish. To determine if the radiance of ocular sparks is equal in all directions, we collected angle-resolved measurements by securing whole triplefins, previously sacrificed by severing the spinal cord, in the center of a platform in a stainless-steel hemisphere of 15 cm diameter placed on a PVC ring holder inside a 7 | Plexiglass^®^ cylinder filled with fresh marine Ringer-solution. The reflective chromatophore patch responsible for generating the sparks was positioned at the exact center of the hemisphere, which was also the exact center of the cylinder, allowing measurements normal to the cylinder wall at all angles. Sparks were generated by means of a stage lamp (ARRI^®^ 650 Plus) mounted ~1.5 m above the fish. To avoid ambient light effects, the room was kept dark. For each of 12 fish, the radiance of the ocular spark was measured with a PR-740 fitted with an MLH-10X lens (Computar^®^) at each 10° between 10° (anteriorly) and 150° (posteriorly) in relation to the frontal-caudal axis of the fish’s body (**Fig. 2**). These values were expressed relative to the radiances of a PTFE diffuse white standard measured at the same angles and position immediately after each fish. The range of angles was not covered for all fish explaining why the sample size varied between angles (**Fig. 2**). Because the lens’ resting state following death is slightly retracted, these measurements could only be used for comparing relative radiance at various angles, but not for estimating relative spark radiance in relation to the illuminant. The size of the ocular spark (*n* = 10) and pupil (*n* = 35) were determined on live fish in a small chamber from scaled images analysed using ImageJ^28^. The light conditions in this small chamber was the same as that described above.

**Figure 2.**
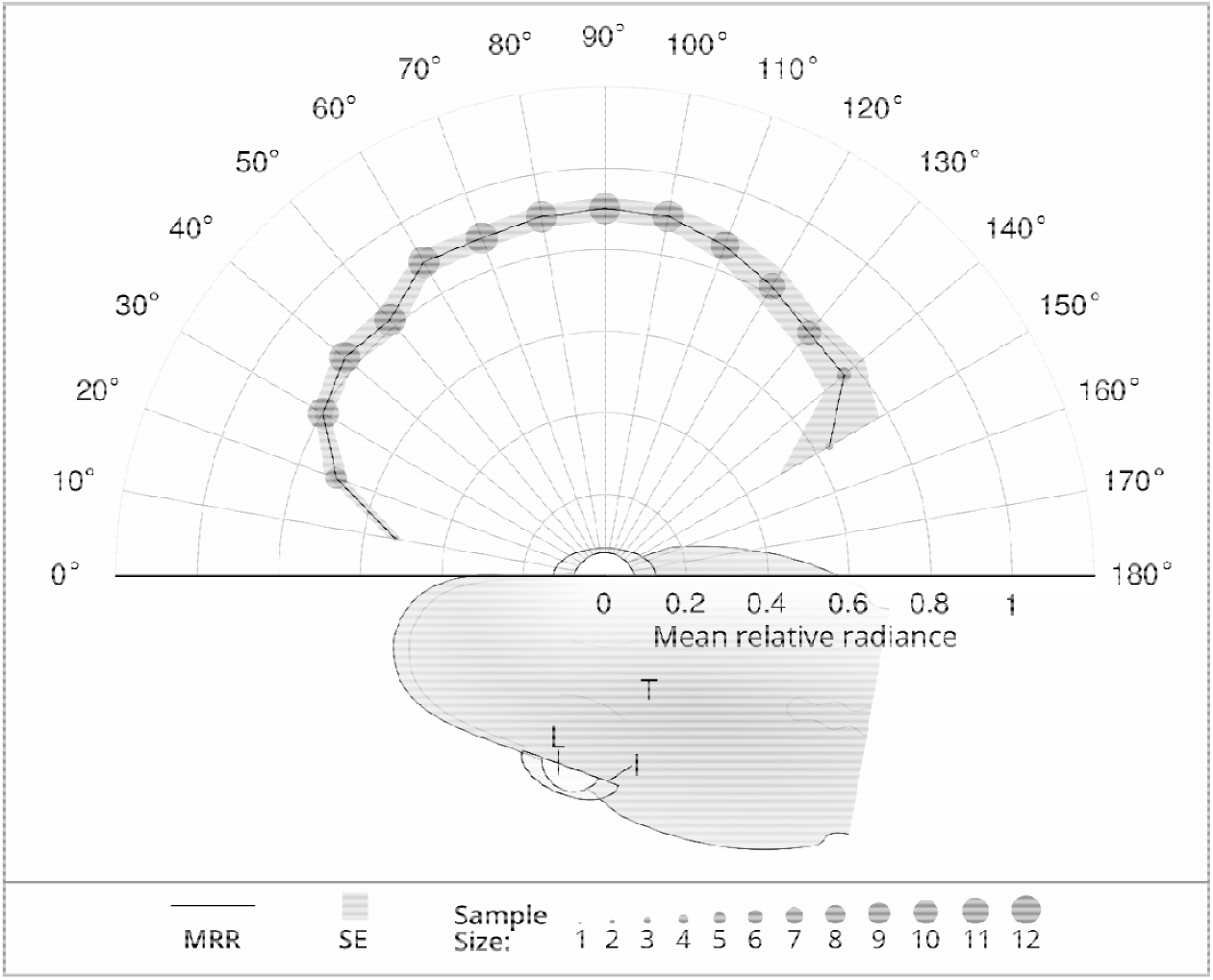
The mean relative radiance (MRR) of the blue ocular spark does not vary along the equatorial axis. MRR is represented by the solid line, the corresponding standard error (SE) by the light grey area, and the sample size of each measured angle by the size of the discs. Fish is seen from a dorsal perspective and naming scheme for angles in relation to the iris of *Tripterygion delaisi:* 90° = normal angle, 0° = angle parallel at the anterior start of the semi-circle and 180° at the posterior end. T = *T. delaisi* body, I = Iris, L = Lens.

### Active photolocation of the gammarid eye

We modeled three-dimensional interactions between triplefins and gammarids, assuming they were both on the same horizontal plane, and that their eyes were positioned at normal incidence. We calculated the photon flux of the reflective eye of the gammarid, as perceived by the triplefin, with and without the contribution of the blue ocular spark, by describing the interaction in simple equations (see complete calculation details in SI). A detectable change in gammarid eye radiance, by switching the ocular spark on and off or through movement of the gammarid eye, would help the triplefin detect potential prey items. In short, the photon flux of the gammarid eye without the contribution of the spark reaching the triplefin retina was determined by the sidewelling light reflected by the ocular reflectors (non-coaxial), the solid angle subtended by the gammarid eye (in steradians) as perceived by the triplefin as a function of the distance between the two eyes, and the area of the triplefin pupil as the ultimate receptor area. The photon flux due to the ocular spark returned to the triplefin was further determined by the radiance of the ocular spark, the solid angle of the ocular spark (in steradians) from the perspective of the gammarid eye, and the coaxial reflective properties of the gammarid’s ocular reflectors. Because solid angle calculations are only possible when the receiver is defined as a infinitely small point in space, not a disc as the pupil of fish or eye of invertebrates, all solid angles were estimated using Monte Carlo simulations^29^.

In each model iteration, we used fixed mean values for parameters that had little influence on the results, based on preliminary sensitivity analyses. We set the gammarid eye radius at 0.0625 mm, the triplefin pupil radius at 0.78 mm, used the downwelling light profile measured at 10 m, the mean background substrate reflectance (*Halopteris filicina*), and the mean reflectance and transmittance of the gammarid body. We explored the parameter space of the possible prey-predator interactions by varying four factors that were determined to have the most influence on the contrast generated by the ocular spark in the gammarid eye (further details in SI).

1. **Spark size:** Because the photon flux that reaches the gammarid eye is directly related to the solid angle subtended by the ocular spark, we varied its radius equivalent on a continuous scale from 0.09 to 0.25 mm.
2. **Spark relative radiance:** The photon flux reaching the gammarid eye is also proportional to the relative radiance of the ocular spark so we varied it on a continuous scale from a mean area under the 400 to 700 nm curve of 0.63 to 2.09.
3. **Gammarid eye reflectance:** The overall radiance of the gammarid eye results from the combined reflection of coaxial and non-coaxial illumination. The non-coaxial component is used to estimate how bright the eye is under the prevailing conditions, without the addition of an ocular spark; the coaxial reflectance is used to calculate the additive contribution of light coming from the ocular spark. We evaluated the impact of the relationship between the coaxial and non-coaxial reflectance of gammarid ocular reflectors using three categories: large difference (non-coaxial reflectance is 9.87 times weaker than coaxial reflectance; maximum observed), average difference (4.09 times weaker), and small difference (2.68 times weaker; minimum observed).
4. **Shading of prey:** Finally, redirecting downwelling light into the horizontal plane would allow triplefins to generate greater contrasts with greater shading of prey, while the triplefin remains exposed to the same downwelling light. We investigated the influence of prey shading using four categories: no shade, weakly shaded, average shade, and strongly shaded. The ‘no shade’ was calculated as the average of the non-shaded sidewelling light measurements divided by the average downwelling light, and the three shaded categories were calculated as the minimum, average, and maximum observed shaded sidewelling light measurements divided by the average downwelling light across the depth gradient described above.

For each set of conditions, we calculated the maximum distance over which active photolocation can contribute to prey detection by calculating the chromatic and achromatic contrast between the gammarid eye with and without the radiance induced by a blue ocular spark as perceived by the triplefin at different distances (range 0.5 – 4.5 cm), and by comparing these values with specific chromatic and achromatic contrast thresholds. The range of distances used is relevant to triplefin feeding behaviour; our own preliminary data from triplefin strikes at prey video-recorded in the wild suggest an average strike distance of 1.31 cm (SD = 0.83 cm, 0.1 – 0.9 quantiles = 0.5 – 2.18 cm, *n* = 107 strikes; Neiße and Michiels unpublished data). Furthermore, the spatial resolution of *T. delaisi* is conservatively estimated at 6 cycles/degree^30^ which means that the average gammarid eye diameter (0.125 mm) becomes a point source at ~48 mm. To avoid modelling situations in which the gammarid eye is smaller than the smallest detectable point in space by a triplefin, we limited the distance between the triplefin and the gammarid to a maximum of 45 mm. The minimum distance modelled relied on estimates of the distance of nearest focus (~5 mm; based on calculations in ^30^).

### Calculation of chromatic and achromatic contrasts

Using retinal quantum catch estimations based on calculated photon flux, we calculated the chromatic and achromatic contrasts between the radiance of the gammarid eye with (photon flux_1_) and without (photon flux_2_) the contribution of the ocular spark radiance. For chromatic contrast calculations we used the receptor-noise limited model^31^ parameterized using triplefin-specific visual characteristics^20,30^. In short, we used species-specific ocular media transmission values, photoreceptor sensitivity curves based on the single cone (peak at 468 nm), and the double cone (peaks at 517 and 530 nm) following a vertebrate photoreceptor-template^32^, and a relative photoreceptor density of single to double cones set at 1:4 as found in the triplefin fovea^30^. Since the Weber fraction (ω) for colour contrast is not known for fish, we used a value of 0.05 as in previous studies from other groups^33,34^. Chromatic contrast calculations result in measures of just-noticeable differences (JNDs), where values above one are considered to represent the minimum discernable differences between the quantum catches. We calculated the Michelson achromatic contrast as (Q_1_ - Q_2_)/(Q_1_ + Q_2_), where Q_1_ and Q_2_ are the quantum catches of the two members of the double cones which are associated with the achromatic channel, under photon flux_1_ and photon flux_2_ respectively. We used two different achromatic contrast threshold values: an optimistic value of 0.008, which was empirically demonstrated in *T. delaisi* using an optokinetic reflex paradigm^35^, followed by conservative calculations using 0.024^36^. The conservative value used is similar to that found in *Carassius auratus*^37^, *Scardinius erythrophthalmus*^38^, *Gadus morhua*^39^, and *Lepomis machrochirus*^40^. To ensure that the contrasts generated by the ocular spark was only influencing the radiance of the gammarid eye and not the background, we performed the same calculations for the gammarid body.

### Calculation of maximum discernable distance

For each set of model conditions defined in the sections above we determined the maximum discernable distance of the ocular spark radiance reflected by the gammarid eye. This was achieved by calculating the chromatic and achromatic contrasts at each millimeter between 5 and 45 mm per set of conditions, and extracting the first value at which the chromatic contrast was equal to or exceeding 1.0 JND, and achromatic contrast equal or exceeding 0.008 for optimistic models, and equal or exceeding 0.024 for conservative models (**SI Fig. S2**).

### Animal care and permits

Fish were caught at STARESO between 5 and 20 m depth using hand nets while scuba diving in accordance with the station’s general scientific permit. During dives, fish were transported in 50 ml perforated Falcon^™^ tubes (Corning Inc, NY, USA) to permit water exchange. At the field station the fish were held in a 50 L flow-through tank at ambient water temperature, until transferred to facilities at the University of Tübingen, Germany. In these facilities, individuals were kept separately in 15 L flow-through tanks (18°C, salinity 34‰, pH 8.2, 12 L: 12 D light cycle) and fed once per day. The fish were sacrificed under approved permit ‘Mitteilung 29.10.2014’ from the Regierungspräsidium (Referat 35, Konrad-Adenauer-Str. 20, 72072 Tübingen) under the supervision of the animal welfare officer.

### Data archival

All data used in the analyses and preparation of figures will be made available on Dryad upon acceptance, and are available (along with R scripts) to the editors and reviewers during the evaluation process.

## Results

### Spectral properties of Cheirocratus gammarids

Focal-stacking images revealed that the reflective units of gammarid eyes are not found in the optical pathway of the eye (**Fig. 3**), but appear to be between ommatidia akin to those described in *Pullosquilla thomassini, Pseudosquillana richeri*, and *Harpiosquilla sp*.^12^. While these reflectors would not improve vision in dim light such as would *tapetum lucidum* and *stratum argentum*^41^, they would normally help camouflage the gammarid eye by making the otherwise black, photon-absorbing, eye look more like the substrate by reflecting some of the sidewelling light field^12^. Overall eye reflectance, within the 400 to 700 nm wavelength range and illuminated with a coaxial light source (epi-illumination), was on average 9.87, *n* = 18; **Fig 3**). The translucent body of the gammarids transmitted much of the light, making them well camouflaged against most backgrounds (**Fig. 3**). The pooled effects of body transmittance and non-coaxial eye reflectance allows gammarids to reduce the contrast between their eyes and the background on which they sit. However, the reflective properties of the structures between ommatidia could be utilized advantageously by triplefins generating ocular sparks.

**Figure 3.**
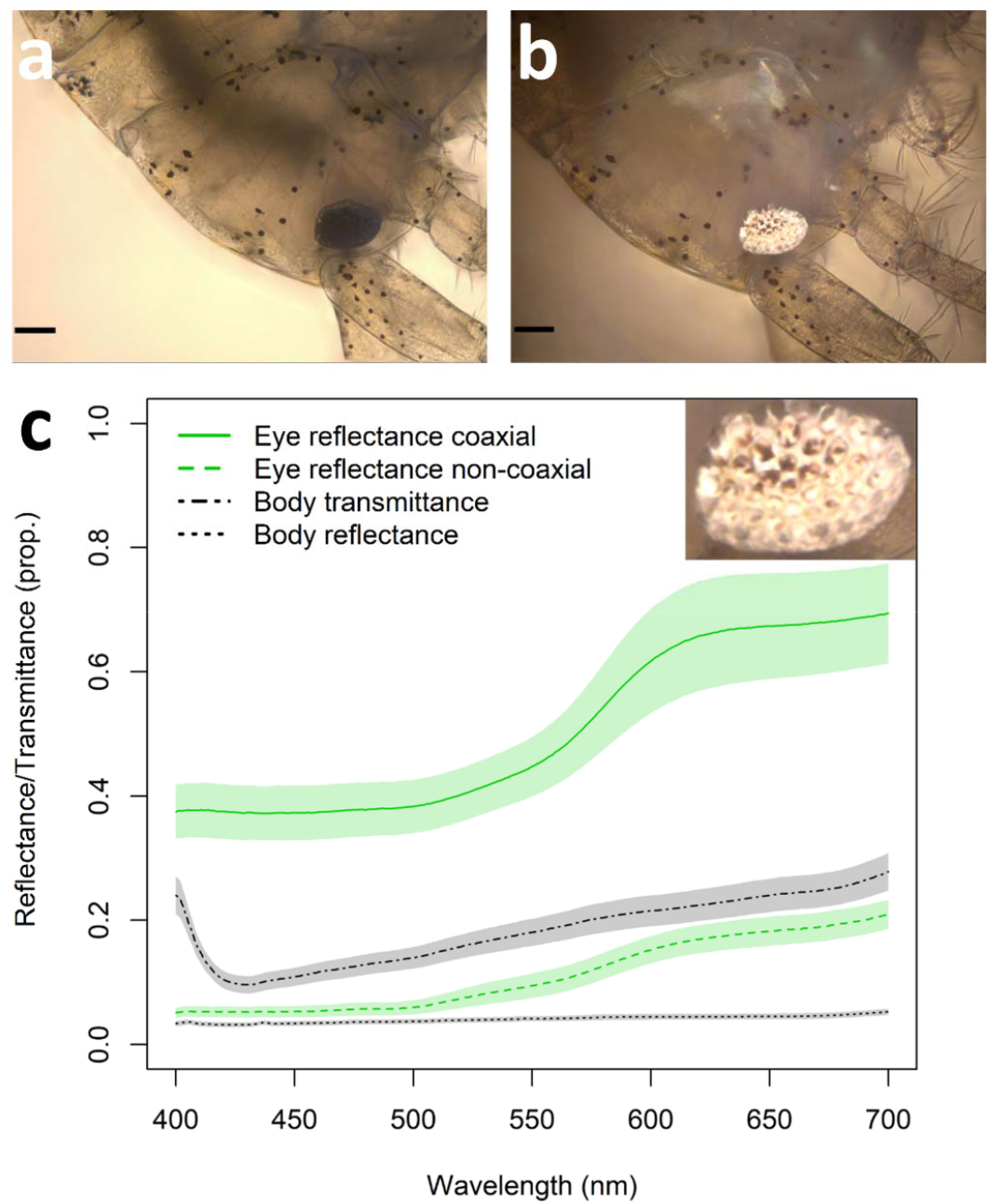
The reflectance of gammarid eyes is much greater under coaxial illumination than non-coaxial illumination. Example gammarid for which measurements of body transmission and reflectance, as well as eye reflectance were obtained; a) viewed under 10 × 10 magnification using transmission illumination, scale bar is 100 μm; b) viewed with coaxial illumination, scale bar is 100 μm. c) Reflectance and transmittance of the body (*n* = 19 individuals) and eye (*n* = 18 for coaxial and *n* = 10 for non-coaxial reflectance); lines indicate average of measurements, shaded area indicate standard error of the mean. Inset shows that the highly reflective structures are between ommatidia. Photo credits: Pierre-Paul Bitton.

### Triplefin and gammarid eye size

From scaled pictures, we determined that triplefin pupil size averaged 0.78 mm (range 0.66 mm to 0.92 mm, *n* = 35 fish, one eye each) and the gammarid eye size averaged 0.063 mm (range 0.021 to 0.102 mm, *n* = 11).

### Properties of ocular sparks

Previous work showed that the relative radiance of the average ocular spark peaks at wavelengths around 472 nm at 2.15 times that of a diffuse white standard, and that the total area under the curve between 400 and 700 nm averaged 1.34 times that of a Lambertian white standard (range 0.63 to 2.09, *n* = both eyes of 5 fish; data from ^8^). The radius equivalent (area as a circular disk) of the ocular sparks ranged from 0.10 mm to 0.24 mm (mean = 0.16 mm, *n* = 10 fish). The relative radiance of the spark was similar across all angles measured along the equatorial axis (**Fig. 2**).

### Active photolocation of the gammarid eye

Modelling results for optimistic calculations show that diurnal active photolocation would assist with micro-prey foraging under wide ranging conditions (**Fig. 4**) by generating perceivable achromatic contrasts (Michelson contrast higher than 0.008) in the eye of gammarids when modulating the ocular spark. A conservative Michelson contrast of 0.024 limited the parameter space under which active photolocation based on achromatic contrasts would be beneficial (**SI Fig. S1**), but demonstrated nonetheless the potential for ocular sparks to enhance prey detection at distances relevant to triplefin foraging behaviour. Chromatic contrast calculations yielded maximum values of 11 mm under only the most supportive conditions and are therefore considered ineffective for gammarid detection using blue ocular sparks (results not shown). Neither achromatic nor chromatic contrast calculations created perceivable contrasts on gammarid bodies (no detection distance above five mm, results not shown).

**Figure 4.**
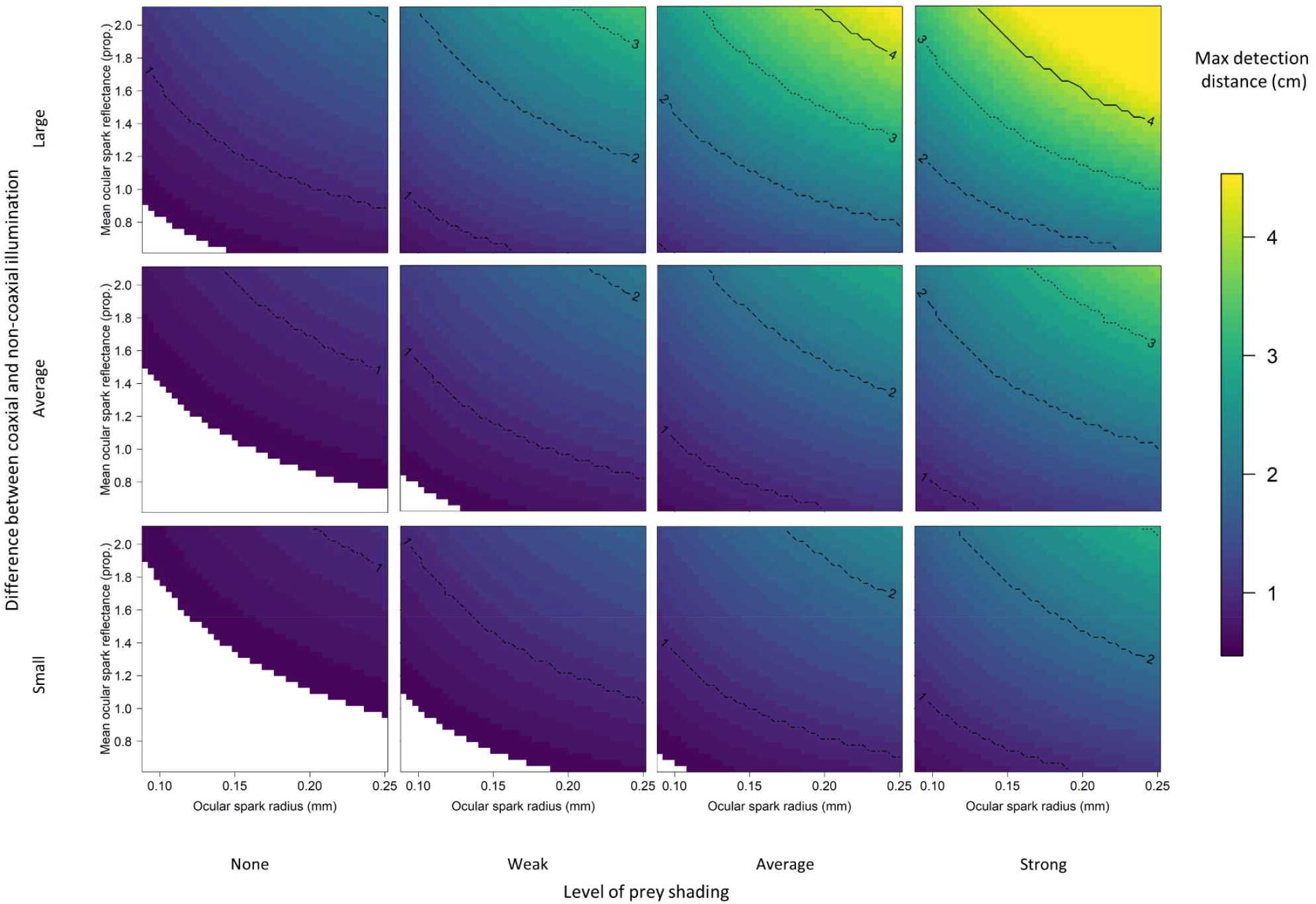
Maximum detection distances of the eye of gammarids by means of blue ocular spark reflectance by triplefins under varying scenarios with Michelson contrast sensitivity set at 0.008. Top, middle, and bottom rows were obtained by varying the relationship between the reflectance of gammarid eyes with coaxial epi-illumination and at 45° from normal. Whereas triplefins were always in the sun, gammarids were tested under four scenarios of shading (columns) in which the prey item is located (See Material and Methods). Conditions in which active photolocation would not assist in gammarid detection are in white.

Under the most favourable conditions, the ocular spark could generate detectable achromatic contrasts at 45 mm distance, the maximum modelled. This distance represents almost a full body length of an average sized triplefin ^42^ and is much longer than their average striking distance (13.1 mm, Neiße and Michiels unpublished data). Under unfavourable parameter combinations, diurnal active photolocation would generate perceivable achromatic contrasts at less than 10 mm, limiting its potential to increase prey detection. In general, diurnal active photolocation would not be beneficial when triplefins forage on unshaded substrates (**Fig. 4** No shade). Under this scenario, only large and bright ocular sparks, and strong coaxial reflectance of gammarid eyes, would generate perceivable achromatic contrasts at distances greater than 10 mm. Even in poorly shaded areas, however, the ocular spark would generate perceivable contrast in the eye of gammarids at greater than 10 mm. When foraging on average or heavily-shaded substrate (**Fig. 4** third and fourth column), the distance at which active photolocation would be beneficial would greatly depend on the relationship between the coaxial and non-coaxial reflectance properties of the gammarid eyes. Under these shaded conditions, maximum detectable distances of over 15 mm would be common, suggesting diurnal active photolocation is effective in many situations.

## Discussion

Through detailed characterization of the interaction involved in the diurnal active photolocation of gammarid prey by *T. delaisi*, we have shown that the controlled production of blue ocular sparks could assist triplefin foraging under a broad range of environmental conditions. It is important to note that active photolocation as described here is a detection enhancement mechanism, not a replacement for regular vision. A gammarid moving through the water column at short distances could obviously be detected through regular vision without any help from active photolocation. However, in the more usual situation where gammarids are well camouflaged because they are translucent and have reflective properties similar to brown algae (see Harant et al.^19^ for algae reflectance), active photolocation could help locate the eyes of previously undetected individuals. In addition, there are many objects on the substrate - as there are in water - that may look like food items, but are not. Hence, even if triplefins detect something that looks like prey, they could decide to investigate it further before striking, maximizing their success. As such, our results are the first theoretical evidence supporting the use of diurnal light redirection in fish as a mean of improving the probability of micro-prey capture.

Furthermore, this study brings into focus the conditions that will facilitate diurnal active photolocation to increase the probability of prey detection. In short, these are: (1) the level of target shading, (2) a means by which the sender/receiver can effectively and coaxially redirect downwelling light, (3) a directionally reflective or retroreflective target, (4) short interaction distances, and (5) sufficient contrast sensitivity in the receiver. We discuss these conditions in details in the following sections.

### Light fields

Our modelling results determined that relatively small differences between the redirected downwelling light and the light field illuminating the gammarid would limit the distance at which ocular sparks would produce detectable contrasts in the eyes of gammarid. Indeed, all three leftmost panels in **Fig. 4** and **Fig. S1** assume that both triplefin and gammarid are directly exposed to downwelling light and show maximum detection distances of perceivable contrast no more than 1.5 cm. This may not seem like a great distance, but preliminary data on foraging triplefins indicate that they normally strike at prey from 1.31 mm on average (Niklas and Michiels, unpublished data). Furthermore, when prey are found in shade the detection distances become much greater (remaining panels in **Fig. 4 and SI Fig. S1**), highlighting the importance of a redirected light field greater than the one illuminating the target for ocular sparks to be effective. Three complementary aspects of the ecology of triplefins and gammarids increase the probability that these favourable interaction conditions are common. First, as in all aquatic environments, the refractive power of the water-air interface constrains downwelling sunrays to a 96° cone, known as Snell’s window, pointing down from the surface^43^. The immediate consequence is that this causes a strong difference in radiance between downwelling and sidewelling light. Second, the environment where triplefins live is very complex and 3-dimensional, generating gradients of light and shade very easily. Indeed, they are often found feeding at small micro-habitat structures such as complex algal growth and encrusting epi-growth^18^. Third, in general, gammarids are substrate-dwelling and only rarely swim in open water, further preferring vegetation with fissured surfaces over those with smooth surfaces^44,45^. Therefore, it is certainly not uncommon for triplefins exposed to direct downwelling irradiance to forage for gammarid that are shaded to a certain degree.

Various factors not considered by our models will undeniably influence the effectiveness of diurnal active photolocation. For example, small waves and wind-generated ripples on the water surface create strong spatio-temporal variation in the downwelling light field near the surface^46,47^. This is observable as highly dynamic light patterns on the substrate, which can vary the irradiance intensity by an order of magnitude at scales smaller than 1 cm, with very short time intervals (milliseconds). These stochastic flashes of light could overwhelm the small differences in reflected eye radiance caused by active photolocation, thus precluding their detection by triplefins. This would be particularly true unless the contrasts generated in the gammarid eye by active photolocation varied asynchronously with the dynamic light pattern. However, both the intensity and frequency of these fluctuations are strongly attenuated as they travel through the water column^47^, and gradually disappear along the 5 – 30 m depth gradiant. Therefore, triplefins foraging between 5 m to 30 m depth would be minimally affected by water surface movement. Of note, however, is that the role for diurnal active photolocation through ocular sparks will also decrease with increasing depth because the ratio between downwelling and sidewelling irradiance decreases due to scattering. Our study does not allow us to determine at what depth diurnal active photolocation through ocular sparks ceases to be effective, mainly because the local substrate had more influence on this ratio than depth. Indeed, the reflective quality of the local vegetative and rocky substrates (pale vs dark) can strongly affect the irradiance of any particular benthic location. The effect of dynamic light field, depth, and local environment need to be further determined to properly determine the importance of these factors in increasing or decreasing the efficiency of diurnal active photolocation.

### Light redirection

The ocular spark was efficient at redirecting the stronger downwelling light field into the weaker sidewelling light field. The average radiance of the sparks was greater than that of a white Lambertian reflector, but there was no evidence that the radiance of the spark varies across the range of angles measured on the equatorial axis of the body plan (**Fig. 2**). While a narrow beam of energy would increase the maximum distance of an active sensing signal, it would also limit the active space from which animals can gather information, leaving them ‘blind’ in other directions^48^. Hence, directional emission would not be particularly advantageous in an active visual sensing system, as the exact position of the reflector would have to be known. Under these circumstances, a broad active sensing signal would be useful for scanning a large area of the visual environment for strong directional reflectors. The fact that ocular spark values radiance is similar across all angles suggests that that the reflective chromatophores have diffuse, not specular, properties. However, ocular spark values greater than that of a diffuse standard demonstrates that downwelling light is being focused onto an area smaller than the lens catchment area. By treating the effect of the lens and reflective chromatophores conjointly, it was simple to determine the emission of the spark with knowledge of the downwelling light field. Further characterization of the spark radiance across the range of angles on the vertical axis will better describe the distribution of the light field redirected by the chromatophores.

The size of the ocular spark had a large effect on the model, simply because the amount of light striking the gammarid eye is strongly dependent on the perceived size of the spark from the gammarids’ perspective. However, producing larger sparks may not be possible or beneficial. The evolution of the size of the reflecting chromatophore patch on which the spark is focused, and therefore the photon radiance available for active photolocation, is probably constrained by two factors. First, the maximum amount of light that can be directed towards the chromatophore is limited by the catchment area of the lens, which depends on the size, position and degree of protrusion through the pupil. This positioning is likely to be driven much more by regular vision than active photolocation. Second, *T. delaisi* is a crypto- benthic species which has evolved colour patterns particularly well suited for camouflage. Generating a large, highly visible spark could become a disadvantage if it attracted potential predators. Indeed, larger piscivorous fish are known to be attracted to bright lures^49^ and several such species are common in the same habitat (e.g. family Serranidae).

Ocular sparks can theoretically be produced by most fishes since a protruding lens is a general feature across teleosts^50^. To produce a functional spark, however, also requires a highly reflective integument where the focussed light strikes the iris. This would occur in species which display a highly localized reflective chromatophore patch, such as in *T. delaisi*, or generally highly reflective irides. In effect, ocular sparks are not the only mechanism by which species could horizontally redirect downwelling light. Completely silvery irides such as those found in zebrafish (*Danio rerio*), sticklebacks (family Gasterosteidae), and a large number of other species could redirect light without the need for focussing. The downwelling light would be reflected by the complete iris, which would redirect a large amount of light almost perfectly coaxially to the pupil. A quick survey of fish eyes under natural conditions does in fact show that light redirecting mechanisms in teleosts are common (see Figure 1 in Michiels et al.^8^), indicating that diurnal active photolocation could be widespread.

### Reflective targets

The difference between the coaxial and non-coaxial reflective properties of the inter-ommatidial structures of the gammarids eyes played an important role in determining the range of distances over which diurnal active photolocation was deemed possible. These reflectors, as those of stomatopod larvae^12^, behaved like imperfect mirrors showing some degree of specularity and diffuse reflectance. Under normal circumstances (i.e., vision by an animal that does not generate or redirect light from structures near its eyes) some of the sidewelling light coming from the animal’s surroundings would be reflected by the invertebrate’s dark eye thus reducing its contrast against the body. However, for triplefins and other fish possessing a light source near their pupil, the viewing geometry between the light source and the receiver (near-coaxial) would make the light reflection stronger.

The presence of reflecting units in the eyes of aquatic invertebrates have been demonstrated in a number of taxa^24-27^, indicating that the benefits of diurnal active photolocation would not be limited to the increased probability of detecting a limited diversity of micro-prey items. While diurnal active photolocation would of course not benefit foraging in detritivores and herbivores, it could offer a substantial advantage to many predators of micro-prey found both on substrate and free floating in the water column. Inter-ommatidial reflectors are not the only mirrors found in nature^11^.

### Interaction distances

The radiative energy emitted in active systems is returned to the sender with an inverse fourth power decrease in light with increasing target distance^48^. As a consequence, any doubling of distance imposes a 16-fold reduction in photon flux. As such, diurnal active photolocation can only be efficient in the context of short distance interactions measured in mm or cm. This implies, as indicated above, that only fish predators of micro-prey are expected to gain foraging advantages through active photolocation. The small scale at which these interactions occur may also help explain why this mechanism has been overlooked until now.

### Contrast detection

Active photolocation cannot work if the animal redirecting and receiving the reflection of light does not have the ability to detect the changes in radiance of the reflecting structure. The detection distance calculations implemented in this study used spatial contrast sensitivity values. However, because we argue that eyes, both those of vertebrates and invertebrates, are likely targets of active photolocation, and because the radiance contrast would be generated by the on/off control of sparks or by gammarids moving their eyes, temporal contrast sensitivity values would be much more appropriate. However, studies of temporal contrast sensitivity are uncommon, especially in non-humans. The limited results indicate that contrast detection of a flickering point in space is influenced, among others, by the size of the stimulus, the amount of ambient light, and the continuity of the field and surround^51,52^. In humans^51,53,54^ as in goldfish^55^ (*Carassius auratus*), the flicker frequency is also an important factor, with contrast sensitivity peaking at around 5-10 Hz in humans but ~ 2-5 Hz in goldfish. At these frequencies, the temporal contrast sensitivity is very similar to the spatial contrast sensitivity, for which data is available from more species, including *T. delaisi*^35^. Furthermore, when attention is paid to the stimulus, temporal contrast sensitivity increases, at least in humans^56^, suggesting the possibility that fish searching for a flickering light caused by their own redirected light could be highly sensitive to a blinking area. In part, this justifies the use of a spatial contrast sensitivity value for assessing a temporal contrast, particularly because we also present results using a conservative threshold of 3 times the measured value. There is no doubt, however, that more research is needed on non-human temporal contrast sensitivity, the factors that may influence it, and its relation with active photolocation.

### Conclusion

Overall, our results describe how active photolocation through blue ocular sparks in the diurnal triplefin *Tripterygion delaisi* could assist in the detection of prey items at relevant foraging distances. We conclude that diurnal active photolocation by means of ocular sparks can supplement regular vision by making the highly reflective eye of potential prey targets shine under nearly-coaxial illumination. Given the high number of fish species that have both protruding lenses and highly reflective irides, active photolocation could be widespread among fish, and an important, yet previously disregarded, vision enhancement mechanism.

## Acknowledgements

We thank the staff at STARESO, especially Corrine Pelaprat for assistance with identification of gammarids, and Oeli Oelkrug for fish maintenance at the University of Tübingen. This project was funded by the German Science Foundation Koselleck grant (Mi 482/13-1) and the Volkswagen Foundation (Az. 89148 and Az. 91816) to N.K.M. as well as running support by the University of Tübingen. P-P.B. was funded by the Natural Sciences and Engineering Research Council of Canada in the form of a Postdoctoral Fellowship (471704 - 2015).

## Author contribution

PPB and NKM conceptualized the study. All authors collected data. PPB, SYC analyzed data. PPB, SYC, and NKM wrote the manuscript, and all authors edited the manuscript.

## Author Declaration

The authors declare no conflict of interest.

