## Supplementary material for "Visual modelling supports the potential for prey detection by means of diurnal active photolocation in a small cryptobenthic fish": SI

Supplementary Information

**Supplementary Table S1.** Symbols used in the equations to calculate the photon flux of the gammarid eye reaching the triplefin, with and without the contribution of the ocular spark

| ***Symbol*** | ***Definition*** |
| --- | --- |
| *L* | Photon radiance (photons s^-1^ sr^-1^ m^-2^) |
| *S* | Blue ocular spark relative radiance (proportion of a PTFE white standard) |
| *d* | Distance between triplefin and gammarid eyes (m) |
| *r*_t_ | Radius of triplefin pupil (m) |
| *R*_ca_ | Reflectance of gammarid eye (coaxial) (proportion of a PTFE white standard) |
| *R*_nca_ | Reflectance of gammarid eye (non-coaxial) (proportion of a PTFE white standard) |
| $\Phi$ | Photon flux coming from the gammarid eye reaching the triplefin pupil (photons s^-1^) |
| *Ω* | Solid angle of target as perceived by receiver (sr) |

Photon flux calculations

We calculated the photon flux of the gammarid eye reaching the triplefin pupil with and without the radiance of the ocular spark, assuming that the center of the triplefin pupil was at normal incidence to the center of the eye of the gammarid, i.e. the full area of the pupil of the triplefin is visible to the gammarid and vice versa. We also assume the effect of absorbance and scattering of the water to be negligible since all energy transfers occur over distances shorter than 5 cm.

*Photon flux without ocular spark*

The base photon radiance of the gammarid eye ($L_{0}$) is a function of the sidewelling light field ($L_{sw}$) and the reflectance of the gammarid eye with non-coaxial illumination:

$L_{0}=L_{sw}\times R_{nca}$	(1)

The photon flux reaching the retina of the triplefin without the ocular spark ($\Phi_{ns}$) is the proportion the gammarid radiance multiplied by the solid angle of the gammarid eye ($\Omega_{gam}$) and the area of the triplefin pupil ($\pi r_{t}^{2}$):

$\Phi_{ns}=L_{0}\times\Omega_{gam}\times\pi r_{t}^{2}$	(2)

*Photon flux produced by ocular spark*

The photon radiance of the ocular spark ($L_{os}$) is a determined by the downwelling light field ($L_{dw}$), the catchment area of the lens, and the reflective properties of the iris chromatophores on which the light is focused. The effect of the lens and reflective properties of the chromatophores have only been measured together and are treated as a relative radiance value (S).

$L_{os}= L_{dw}\times S$	(3)

The radiance of the gammarid eye ($L_{gam}$) caused by the reflection of the ocular spark is estimated by multiplying the radiance of the ocular spark reaching the gammarid ($L_{os}$) with the solid angle of the ocular spark ($\Omega_{os}$) and the reflectance of the gammarid eye with illumination coaxial to the receiver ($R_{ca}$). Because the properties of the gammarid eye are measured in relation to a diffuse white standard, the photon exitance from the gammarid eye is converted to photon radiance by dividing by $\pi$ steradians:

$L_{gam}= L_{os}\times\Omega_{os}\times R_{ca}\times\pi^{-1}$	(4)

The photon flux generated by the ocular spark which reaches the triplefin retina ($\Phi_{os}$) is determined as the proportion of the ocular spark generated gammarid eye radiance (Eq. 4) multiplied by the perceived size of the gammarid eye, in steradians, and the area of the triplefin pupil:

$\Phi_{os}=L_{os}\times\Omega_{os}\times R_{ca}\times\pi^{-1} \times\Omega_{gam}\times{\pi r}_{t}^{2}$	(5)

The total photon flux reaching the retina of the triplefin with the ocular spark is then the sum of equations (2) and (5).

A similar calculation was used for the effect of the ocular spark on the illumination of the gammarid body. In these calculations we estimated the photon flux reaching the retina of the triplefin with and without the contribution of the ocular spark, using the same solid angles. In contrast to calculations with the gammarid eye, we used the same body reflectance values for the coaxially and non-coaxially illuminated scenarios. The photon exitance from the body, both with and without the contribution of the ocular spark was determined as the proportion of light that was reflected by the body and the proportion of light that was transmitted through the body, reflected by the substrate, and transmitted again through the body.

For all calculations, the solid angle of the gammarid eye from the perspective of the triplefin pupil ($\Omega_{gam}$), and the solid angle of the ocular spark from the perspective of the gammaridh eye ($\Omega_{os}$), in steradians, were estimated by Monte Carlo simulation (35). The triplefin pupil, gammarid eye, and ocular spark were treated as disks of zero thickness. The pupil and gammarid eye were always positioned centered and at normal incidence to one another, and the ocular spark positioned at the edge of the iris (displacement = 1.09 mm) in the same plane and normal vector as the triplefin pupil. Because we estimate that the triplefin can focus on objects minimally at 7 mm and that average gammarid eye becomes a point source beyond ~48 mm, we determined the solid angles for distances between 5 mm and 45 mm. The calculations were based on 1E09 photon packets emitted from the source; these generated solid angle estimates with 99.9% confidence intervals with errors ranging from 1.2 % of the solid angle value at 5 mm to 10.6 % at 45 mm.

Exploration of parameter space

To explore the parameter space of our interaction between triplefins and gammarids, we varied the parameters known to have the most influence on the calculated contrasts. To allow comparison and visualization of the results, we chose to model two continuous parameters: the ocular spark radius and the ocular spark relative radiance, and two categorical parameters: the relationship between the coaxial and non-coaxial reflectance of the gammarid eyes, and the relationship between the downwelling and sidewelling light field.

The parameter ‘ocular spark radius’ ranged from 0.09 mm to 0.25 mm (based on actual measurements ranging from 0.10 mm to 0.24 mm) in 41 intervals of equal increments (0.004 mm). The range values for the parameter ‘ocular spark relative radiance’ was produced by first taking the mean value of all measurements at each wavelength (binned in 1 nm interval) and varying the area under the curve between the measured range of 63 % to 209 %. To produce square matrices of results, the value range was also divided in 41 intervals of equal increments.

The relationship between the coaxial and non-coaxial reflectance of gammarid eyes was not correlated in the samples measured. To explore the influence of this parameter we calculated the average difference between the coaxial and non-coaxial eye reflectance measurement obtained from each gammarid, calculated at each wavelength (binned in 1 nm interval), and varied the area under the curve to represent the minimum value observed (10.1 %), the average value (24.4%), and the maximum value observed (37.25 %).

We included four measures of the relationship between the downwelling and sidewelling light fields: no shade, weakly shaded, average shade, and strongly shaded. The relationship between the downwelling and unshaded sidewelling spectral light profile was obtained by taking their ratio at several measured locations. The three categories of shaded sidewelling light were obtained by calculating the average difference between the downwelling and shaded sidewelling light fields at each measurement station, and varying the area under the curve to represent the minimum value observed (ratio DW/SW = 8.65), the average value (ratio = 16.62), and the maximum value observed (ratio = 26.63). These conversion vectors were then applied to the downwelling light field obtained at 10 m depth.

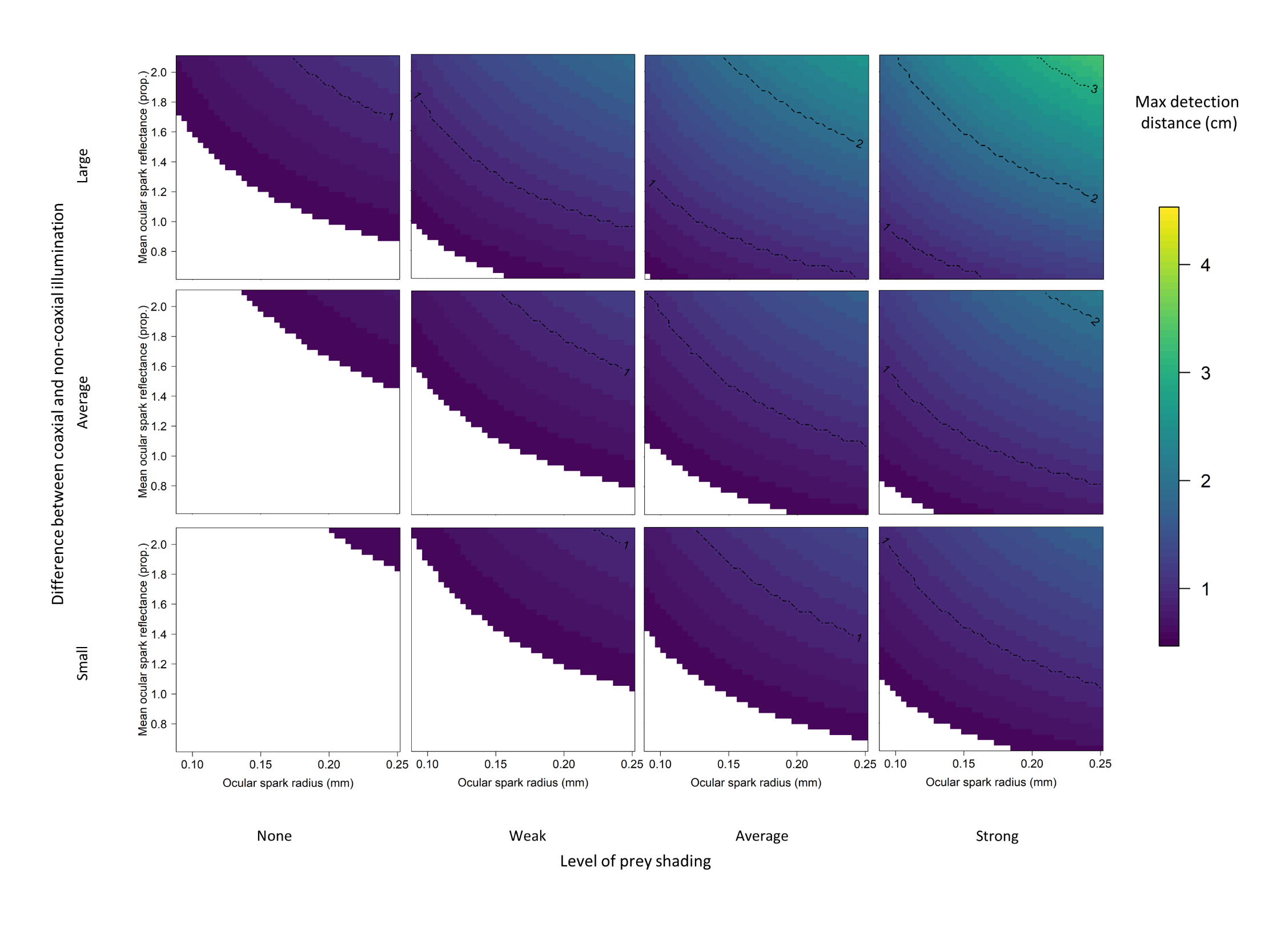

**Supplementary Figure S1.** Maximum detectable distances of ocular spark reflectance from the eye of gammarids under varying scenarios (Weber fraction = 0.05). Top, middle, and bottom row were obtained by varying the relationship between the reflectance of gammarid eyes with coaxial epi-illumination and at 45° from normal. Vertical rows were obtained by varying the amount of shade on which prey items rests. Conditions in which active photolocation would not assist in gammarid detection are in white.

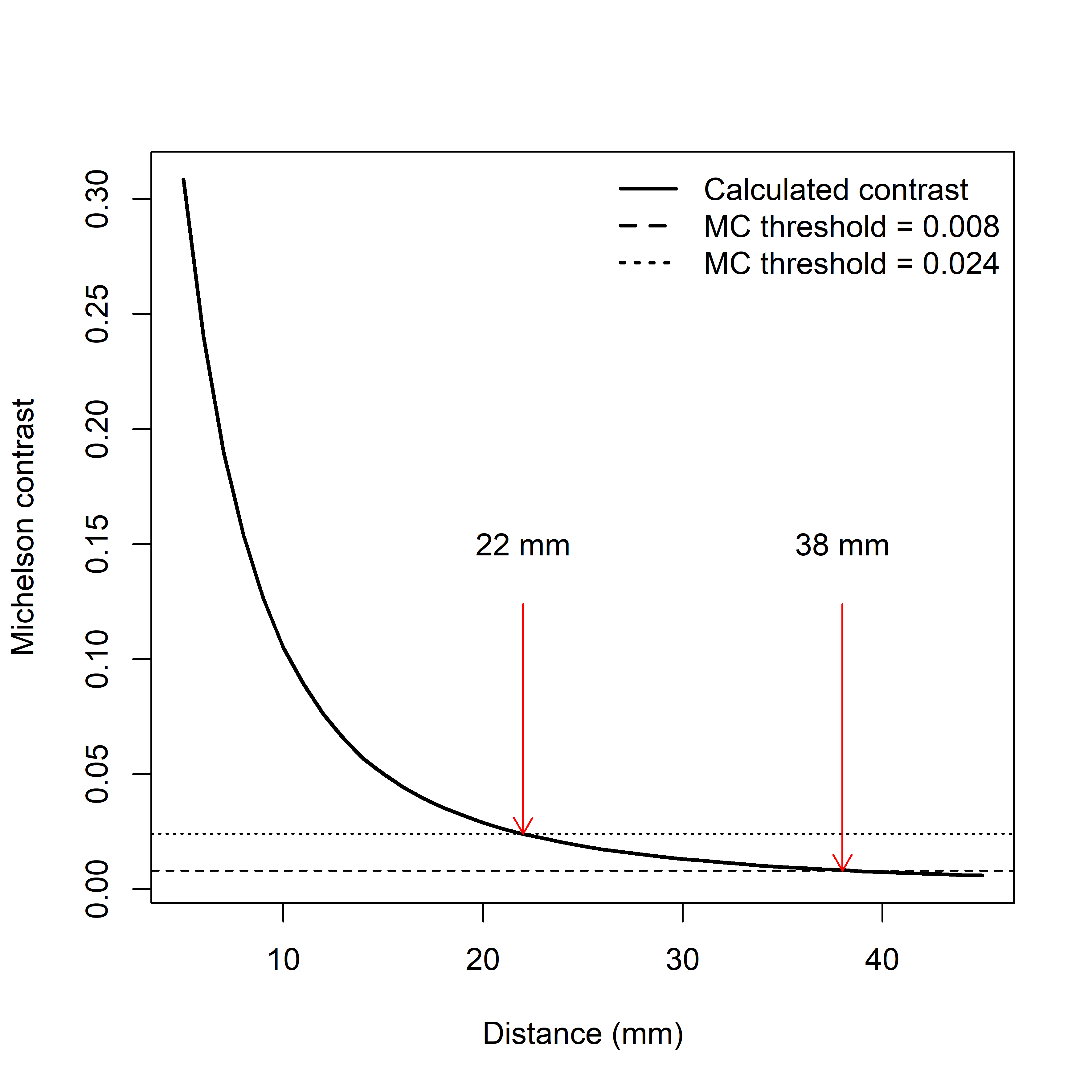

**Supplementary Figure S2.** Example extrapolation of the maximum distance at which reflections in the gammarid eye caused by ocular spark radiance are discernable. The Michelson contrast (achromatic contrast) is the perceived difference in photon flux from the gammarid eye with and without ocular spark contribution. The maximum discernable distance is defined as the distance at which the Michelson contrast is equal to an optimistic value of 0.008, or a conservative value of 0.024.
